# Water-triggered, irreversible conformational change of SARS-CoV-2 main protease on passing from the solid state to aqueous solution

**DOI:** 10.1101/2021.05.21.445090

**Authors:** Narjes Ansari, Valerio Rizzi, Paolo Carloni, Michele Parrinello

**Affiliations:** Italian Institute of Technology, Via E. Melen 83, 16152, Genova, Italy; Computational Biomedicine, Institute for Advanced Simulation (IAS-5) and Institute of Neuroscience and Medicine (INM-9), Forschungszentrum Jülich, Jülich, 52425, Germany, JARA-Institute Molecular Neuroscience and Neuroimaging (INM-11), Forschungszentrum Jülich, Jülich, 52425, Germany. Physics Department, RWTH Aachen University, Aachen, Germany

**Author notes:** Contributed equally to this work (Narjes Ansari, Valerio Rizzi).

## Abstract

The main protease from SARS-CoV-2 is a homodimer. Yet, a recent 0.1 ms long molecular dynamics simulation shows that it readily undergoes a symmetry breaking event on passing from the solid state to the aqueous solution. As a result, the subunits present distinct conformations of the binding pocket. By analysing this long time simulation, here we uncover a previously unrecognised role of water molecules in triggering the transition. Interestingly, each subunit presents a different collection of long-lived water molecules. Enhanced sampling methods performed here, along with machine learning approaches, further establish that the transition to the asymmetric state is essentially irreversible.

---

The appearance of a new coronavirus in December 2019 has triggered a pandemic that is spreading throughout the world^1,2^. The virus is closely related to other coronaviruses^3^ such as SARS-CoV and MERS-CoV and has been named SARS-CoV-2. As a triumph of modern science, a number of vaccines were developed at an unprecedented fast rate and have been provided to a significant fraction of the human population^4^. However, these vaccines might be inefficient against constantly emerging new variants of the virus. Thus, it is imperative to advance drug discovery campaigns to identify ligands which may interfere with the disease^5^.

One of the main drug targets for anti SARS-CoV-2 infection is the main protease (SC-2 M^pro^ hereafter)^6–12^ that specialises in cleaving the nascent viral polypeptide chain. This enzyme is a homodimer and the presence of the two subunits is required for its biological function^12^. Each of the subunits is formed by three domains: domains I (residues 10-99) and II (residues 100-182) have an antiparallel *β*-barrel structure, while domain III (residues 198-303) forms a compact *α*-helical domain connected to domain II by a long linker loop^10^ (Fig. 1 (a)). The enzymatic reaction is performed by the catalytic His41-Cys145 dyad, located in the S2 site of each of the subunits (see Fig. S-1 in Supporting Information (SI)),^13,14^. Intriguingly, although most X-ray studies on SC-2 M^pro^ show an almost perfect symmetry of the homodimer^7^, several indirect lines of evidence suggest that only one of the two subunits is catalytically active in each of the subunits (as seen for a variety of other homodimeric proteins^15^): (i) In the almost identical homodimeric protein from SARS-CoV (SC M^pro^, 96% sequence identity with SC-2 M^pro^)^7^, one subunit is inactive because of distortions of the active site^16–19^; molecular dynamics (MD) provided insight on this asymmetry of the protein^18^, similarly to what has been done for other homodimeric proteins^20,21^. (ii) Different analyses of a 0.1 millisecond-long MD simulation recently performed by D. E. Shaw’s research group (DESRES)^22^ of SC-2 M^pro^ in aqueous solution suggested that each of the two subunits visits a different set of configurations, after starting from a symmetric X-ray structure (PDB 6Y8423)^24–26^. This hints at a role of packing forces for affecting the conformation of the protein^27^. Consistently, other MD studies suggested that only one subunit is catalytically active^28,29^.

Here we re-analyse the DESRES simulation22 focusing on symmetry breaking important factors for the protein. The key role of water is clearly apparent from the very first part of the trajectory in which the initial symmetry is broken. In fact, we observe a single water molecule entering the binding pocket of one of the subunits, which starts a series of changes that lead to the disruption of the catalytic site and to symmetry breaking. In particular, a key part of the observed chain of events, namely the contact breaking between the N-finger terminus (residues 1-7, Fig. 1 (b)) of one subunit and the m-shaped loop (residues 135-146, Fig. 1 (b)) of the other one, was known to play a key role in SC M^pro^ dimerization and (in)activation^17^. After this initial phase, we observe the emergence of a long lived asymmetric state. In this state, the binding pocket in the two subunits have different volumes and contain a different number of water molecules (see SI).

**Figure 1.**
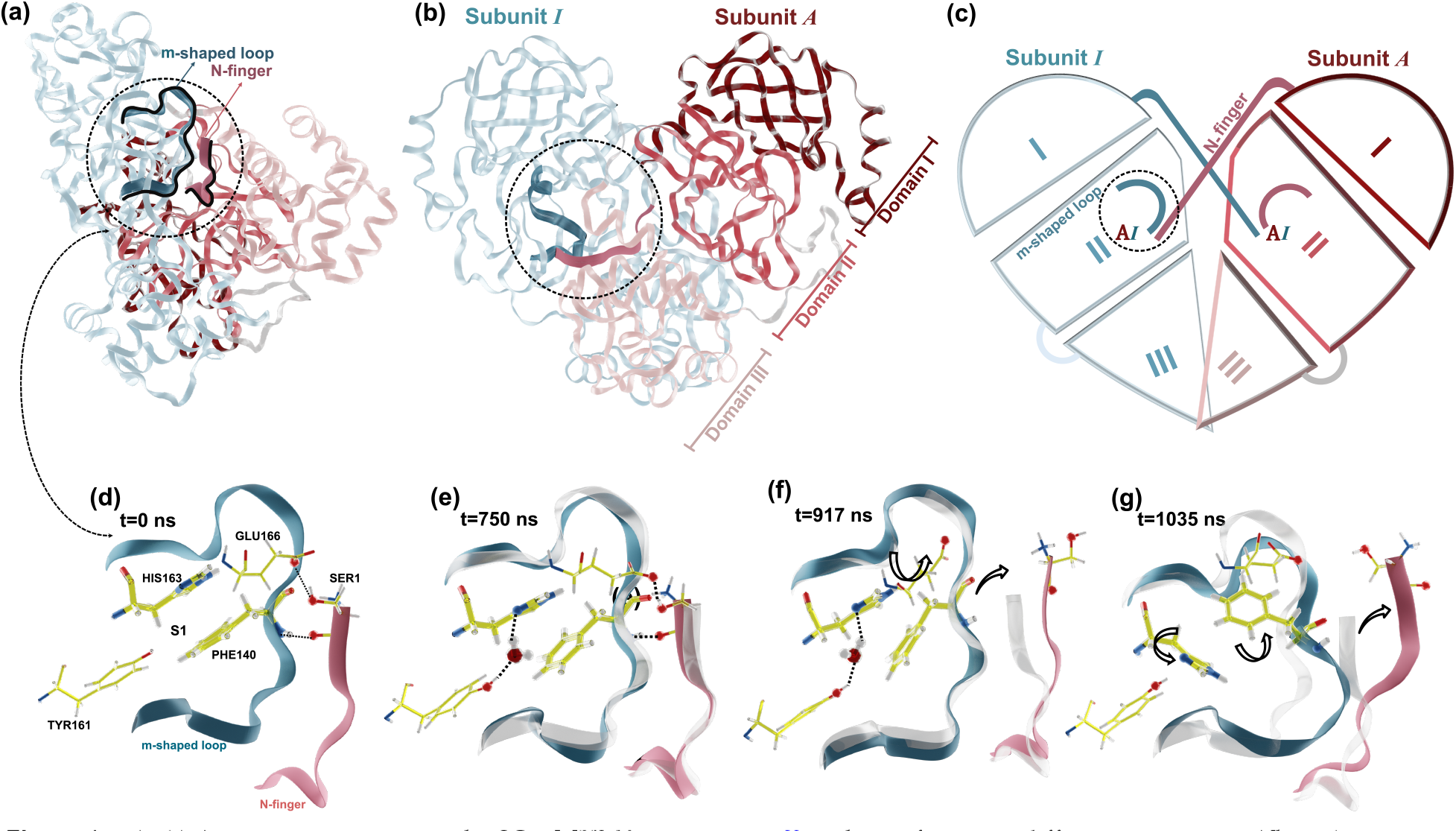
(a, b) A cartoon representing the SC-2 M^pro^ X-ray structure 23 is shown from two different viewpoints. The subunits, ***I*** in light blue and ***A*** in red, consist of domains I-III (discussed in the text). The m-shaped loop (residues 135-146 of ***I*** and the N-finger (residues 1-7 of *A*), discussed in the text, are shown in darker colours and they are enclosed in a dashed circle. (c) Schematic of the protein as shown in (b). (d)-(g) Symmetry breaking of subunit ***I*** on passing from the X-ray structure to water solution, as observed in the DESRES simulation^22^. This event occurs between 0.75 and 1.035 *μ*s. For the sake of simplicity, only residues relevant to the process are shown; (d) the initial conformations of the m-shaped loop and the N-finger conformations (also in (a) and (b)). The two regions bind to each other via the Glu166@***I*** - and Phe140@***I*** - Ser1@***A*** H-bonds. The intra-subunit interaction is supported by His163@***I*** and Phe140@***I****π*-stacking. (e) One water enters the S1 binding pocket and forms a bridge between His163@***I*** and Tyr161@***I***. (f) A rotation of Glu166@***I*** leads to a breakage of its H-bond with the N-finger. (g) A water molecule that leaves the S1 binding pocket triggers a disruption of the hydrophobic contact between residues His163@***I*** and Phe140@***I***, and eventually leads to the inactivation of subunit ***I***.

Could the protein swap from one asymmetric conformation to the other? Our analysis helps us in identifying a slow mode of the system that expresses a subunit active-inactive transition. By amplifying its fluctuations in the symmetric state by means of a newly developed enhanced sampling method^30^, we observe inactivation events in one subunit that follow an analogous path as the one observed above. When starting from the asymmetric state, we are able to induce transformations in which the subunits exchange their role. However, our estimate of the corresponding kinetic barrier is so high that on the biological timescales the subunit flip is unattainable. This reported stability of the homodimer asymmetric state is practically irreversible.

Let us now describe these results in more detail. We start by illustrating how the initial symmetric configuration (PDB entry 6Y84^23^) shown in Fig. 1 (a,b) breaks its symmetry in DESRES simulation^22^. We follow in particular the intra- and inter-subunit contacts which have been shown to play an essential role for the stabilisation of the S1 binding pocket in SC-2 M^pro^. Such contacts are the Phe140/His163 intra-subunit hydrophobic interaction and the inter-subunit interaction between the m-shaped loop and the N-finger, in particular the Glu166/Ser1 H-bond (Fig. 1 (d)). We name the two subunits ***A*** and ***I*** anticipating that eventually the first one will remain active and the other one will become inactive. Following the observations from Ref. 16, the active form, in contrast to the inactive one, satisfies simultaneously two main criteria which are essential in stabilising the S1 binding pocket (Fig. 1 (d)). First, it shows an intra-subunit *π*-stacking contact between side-chains Phe-140 and His-163 (Fig. 1 (d)). Second, it has an intersubunit hydrogen bond between its Glu-166 residue and the Ser-1 residue of the other subunit (Fig. 1 (d)).

After 750 ns, a water molecule bridges the imidazole nitrogen atom of His163@***I*** and the hydroxyl oxygen atom of Tyr161@***I*** (see Fig. 1 (e)). After about 150 ns, Glu166@***I*** rotates. Following that, the N-finger@***A*** changes its conformation, and disrupting the link between the m-shaped loop@***I*** and the N-finger@***A***. Eventually, the water molecule exits from the S1 binding pocket, leaving behind an empty space that probably encourages residues His163@***I*** and Phe140@***I*** to rotate away from one another, breaking the catalytic dyad (see Fig. 1 (g)). The structural determinants of subunit ***I*** is to be contrasted with that of subunit ***A*** where the catalytic dyad remains stable. This symmetry broken state is maintained for the rest of the s1mulation (about 99 *μ*s).

**Figure 2.**
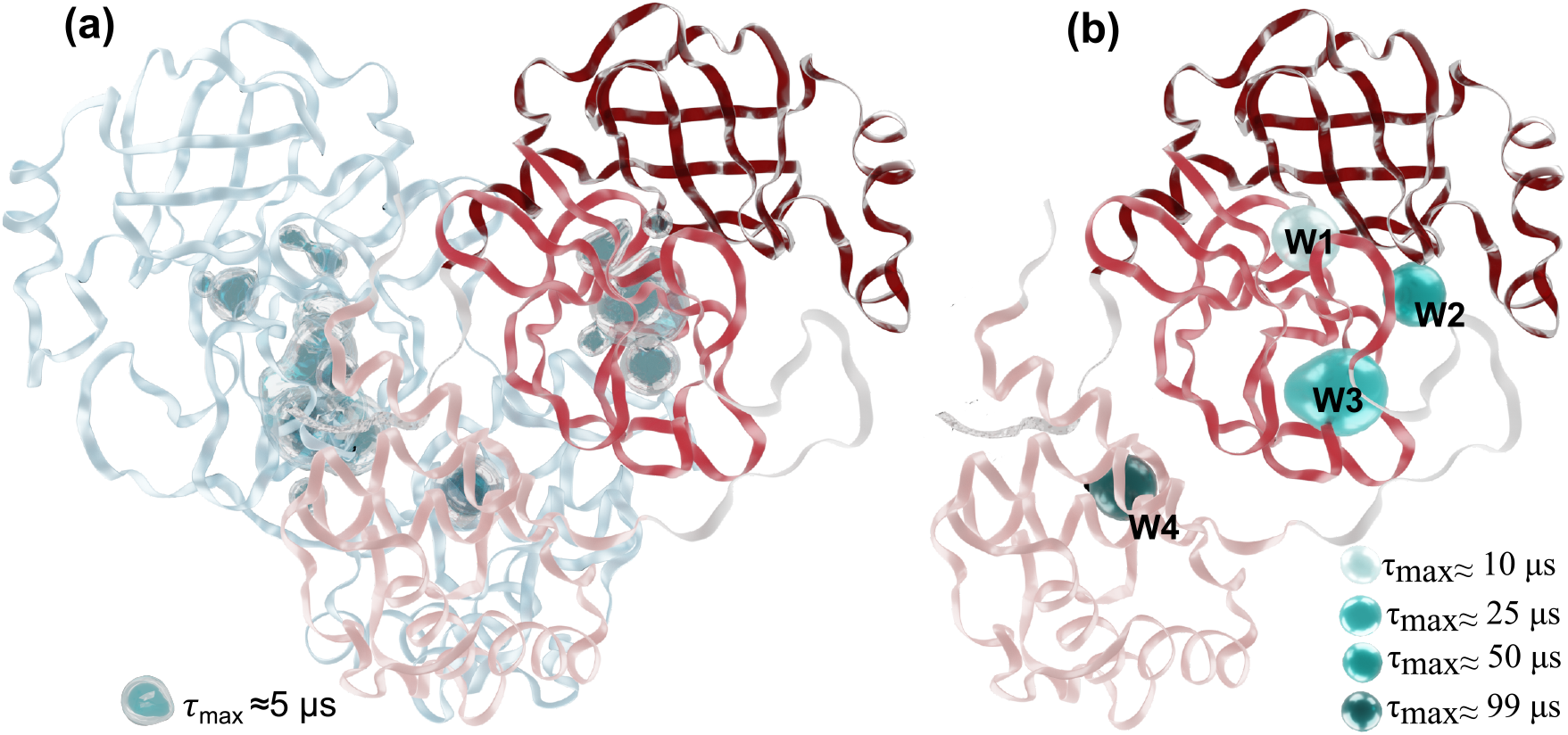
The average position of (a) medium- and (b) long-lived water molecules (for their definition, see text) inside of the two subunits of a SC-2 M^pro^ after 60.2 *μ*s. In the former case, the distribution of medium-lived water molecules is asymmetric and delocalised, while in the latter case, the distribution is symmetrical in both subunits and consists of 4 hydration sites (W_1_-W_4_). The water maximum residency time *τ*_max_ is shown in *μ*s.

We conclude that water (within the well-known force field limitations^31^) does play a role in the disruption of one of the catalytic sites. This prompts us to investigate water dynamics. We allocate the water molecules inside the subunit in two groups according to their lifetime during the last 99 *μ*s.

The *long-lived water molecules* (with a lifetime larger than 5 *μ*s) are located in four spots (W_1_-W_4_) whose position is symmetrical in the two subunits (Fig. 2 (b)). The lifetime increases on passing from W_1_ to W_4_. W_1_ and W_2_ bridge domains I and II, while W_3_ and W_4_ sit in domains II and III, respectively. In W_4_, water is deeply buried and stays for the entire dynamics. If a water molecule leaves W_1_-W_4_, it is quickly replaced by another one, supporting the notion that water in these spots may have a structural role. The three longest-lived spots (W_2_-W_4_) are in close proximity with water molecules from X-ray diffraction data (PDB entry 6Y84). The long lifetime of the W_2_-W_4_ spots is consistent with the very low values of the Debye-Waller factors for these three water molecules (see Tab. S-1 and Fig. S-3 in SI).

The *medium-lived water molecules* (with a lifetime between 0.5 *μ*s and 5 *μ*s) sit preferentially in non-symmetry related positions (see Fig. 2 (a)), thus contributing to the asymmetry of what appears to be the equilibrium state. They occupy a region that mostly encompasses the S1 binding pocket (see Fig. 3). We shall refer to this region as the water reservoir (see SI for further details) and to the positions where water accumulates as hydration sites. The asymmetry between subunits ***I*** and ***A*** is mirrored in the larger number of hydration sites in subunit ***I***. Furthermore, in subunit ***A***, the water reservoir cavity is essentially rigid and has a rather constant water occupation, while in subunit ***I*** it exhibits volume fluctuations correlated with its content of water molecules (see SI).

The way that the two subunits interact and their respective conformation is correlated with their medium-lived water content. In subunit ***I***, the m-shaped loop strongly interacts with the N-finger of subunit ***A*** via several hydrogen bonds, see Fig. 3 (a). As a result of these interactions, the large aromatic ring of Phe140@***I*** flips (Fig. 3 (a)). This event opens up a path for water molecules to enter, leading to the collapse of the S1 pocket (indicated by label (3) in Fig. 3 (a)). Because of the reorientation of the m-shaped loop and the hydrophobic *π*-stacking of His163@***I*** and Phe140@***I***, the S1 binding site of subunit ***I*** collapses. On the other hand, in subunit **A**, the interaction between the m-shaped loop@***A*** and the N-finger@***I*** is different (see Fig. 3 (b)). The Ser1@***I*** amino forms a hydrogen bond with the carbonyl oxygen of Glu166@***A*** and together with the hydrophobic interaction between the His163@***A*** and Phe140@***A*** stabilises the catalytic site (see label (3) of Fig. 3 (b)).

To determine whether the symmetry breaking can be reversed, one can use enhanced sampling simulations^32^. Here, we apply the recently developed on-the-fly probability enhanced sampling (OPES) method^30^ that is the latest and arguably most efficient evolution of Metadynamics^33,34^. As in Metadynamics and in many other enhanced sampling methods, OPES requires the definition of appropriate collective variables (CVs) whose fluctuations are enhanced in a statistical mechanics compliant way^32^. Here, the CV construction is facilitated by the insight gained by our examination of the DESRES trajectory. Our procedure requires two steps. At first, we identify a number of descriptors that are able to distinguish whether the catalytic site is active or inactive. Because of this, we choose 26 descriptors that focus on the region in the vicinity of the m-shaped loop, the N-finger and the S1 binding pocket (see SI for more details), around which much of the action takes place. It is interesting to note that none of these descriptors directly reflect water behaviour. However, in the choice of the descriptors we were guided by the location of the hydration sites and have included in the descriptors information on the local protein conformation. Thus, we might say that water, albeit indirectly, is included in the set of descriptors.

**Figure 3.**
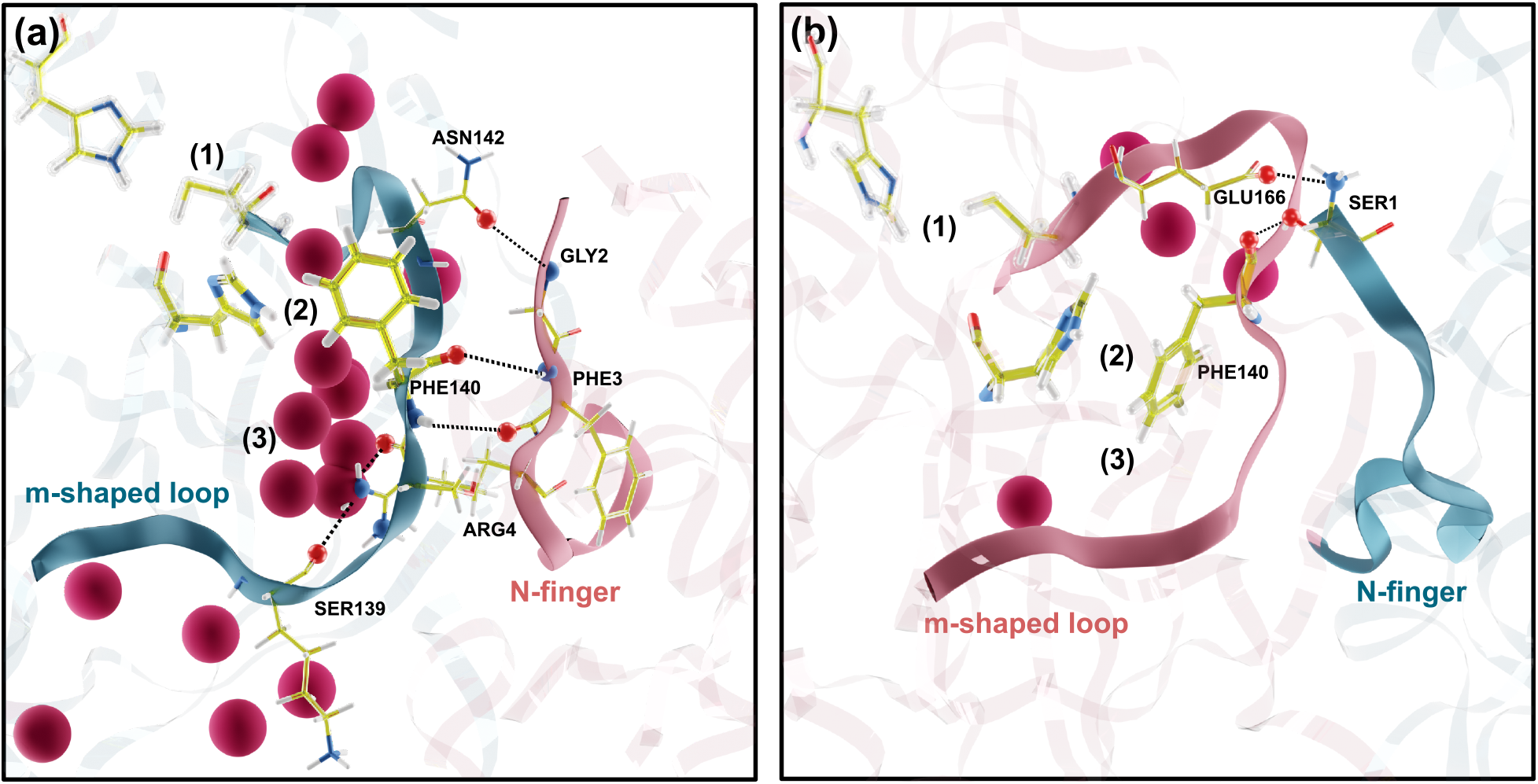
Conformations of the m-shaped loop and the N-finger in (a) the inactive ***I*** and (b) the active **A** subunits in an equilibrated structure of the DESRES simulation (namely, after 60.2 *μ*s). The positions of hydration sites with a water occupancy larger than 50 % of the time are shown as red spheres. The position of the catalytic dyad, the stacking between Phe140 and His163 and the S_1_ binding pocket, are shown as labels (1)-(3), respectively. Relevant hydrogen bond interactions between the two subunits are highlighted by a dashed-line.

As a second step, we apply Deep-LDA^35,36^, a machine learning-based procedure that, given a set of descriptors and their fluctuations, is able to generate efficient CVs in the study of transitions between metastable states. In our case, the states in question are the configurations of the active subunit and the inactive one from the DESRES trajectory. By applying the resulting CV independently to both subunits and starting from the symmetry broken homodimer state, we aim to trigger transitions to the flipped homodimer where the two subunits exchange their role.

Given the complexity of the system and its computational cost, one cannot expect the system to undergo several transitions between one state and another and fully converge a free energy surface. However, we find it encouraging that after 20 ns of an exploratory OPES simulation the system does make a transition where the identity of the subunits is reversed in a rather smooth way without passing via unnatural conformations (see Movie in SI). A rough estimation of the corresponding free energy barrier is possible (see SI). We estimate the barrier to lie between 125 to 150 kJ/mol which corresponds to an extremely rare event at room temperature whose characteristic time which far exceeds the SC-2 M^pro^ lifetime. Of course, such an estimate must be taken with a lot more than a pinch of salt. Nevertheless, we can be reasonably confident to say that the transition of SC-2 M^pro^ into an asymmetric homodimer state is indeed irreversible.

In conclusion, our analysis of the very long simulation in Ref. 22 on SC-2 M^pro^, complemented by enhanced sampling simulations, suggests that water molecules present in one of the two subunits triggers the transition from a symmetric to an asymmetric state, in which only one of the two subunits is functional. Such transition is irreversible.

## Acknowledgement

We acknowledge the Italian Institute of Technology for funding and support. The simulations were performed on the supercomputer CLAIX-2016-GPU at Forschungszentrum Jülich. Part of this work was performed under the auspices of ETH Zürich and Università della Svizzera italiana, Lugano. We thank David Shaw’s group for making their long MD trajectory available to the scientific community.

