## Supplementary Information for "Water-triggered, irreversible conformational change of SARS-CoV-2 main protease on passing from the solid state to aqueous solution"

[∇]Contributed equally to this work

May 21, 2021

#### Contents

**S1 Volume and number of water molecules contained in each subunit** **S2**

**S2 Mobility of the water molecules inside the protein** **S4**
S2.1 Low mobility . . . . . . . . . . S4
S2.2 All ranges of mobility . . . . . . . . . . S5

**S3 Hydration sites** **S6**

**S4 Volume and number of water molecules contained in S1 binding pocket.** **S6**

**S5 Symmetry of the two subunits** **S8**

**S6 Details of the simulations carried out in this work** **S8**

**S7 Plain molecular dynamics** **S8**

**S8 Enhanced sampling simulations** **S8**

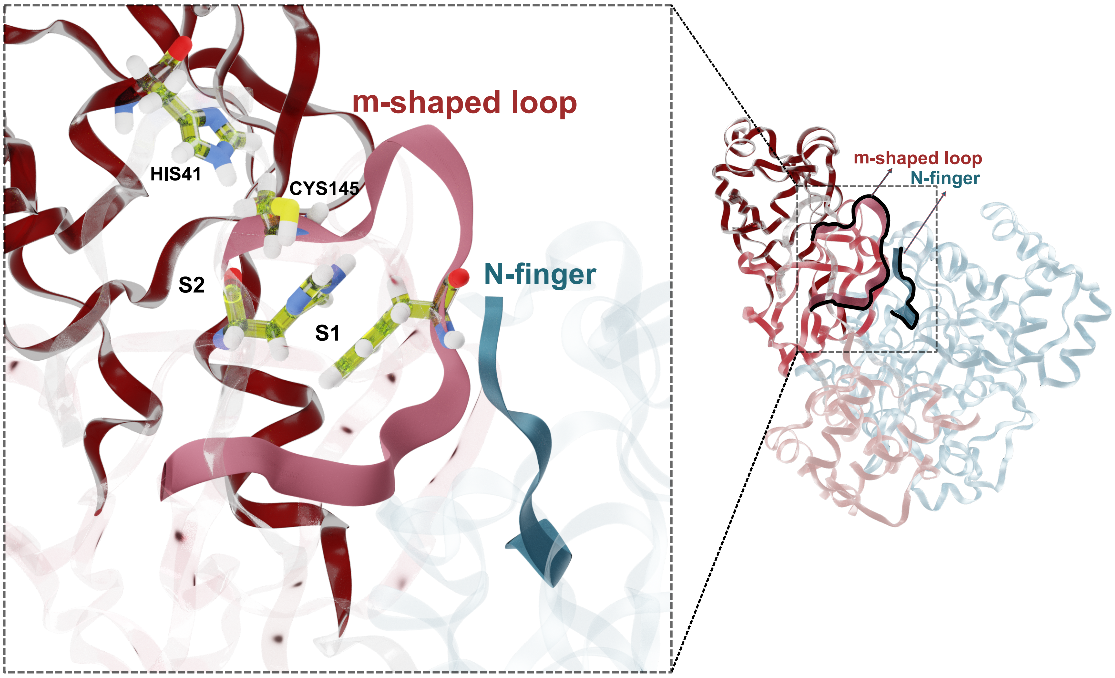


Figure S-1: The catalytic Cys-His dyad in the SC-2 M$^{pro}$ S2 binding site and its location in subunit $\boldsymbol{A}$. The structure represented here is the initial one of the DESRES deposited trajectory [1], that is the X-ray structure (PDB entry 6Y84) [2]) after undergoing MD equilibration.

#### S1 Volume and number of water molecules contained in each subunit

We calculate the volume of each subunit (Fig. S-2 (a-b)) by defining a convex hull surface whose vertices are 112 $\alpha$-Carbon atoms belonging to residues on the subunits' surfaces (see panel (a) of Fig. S-2). The C-terminus tails, highly mobile, are not included. The distribution of the volume and the water content inside the convex hulls in the DESRES trajectories are shown in Fig. S-2 (c) and Fig. S-2 (d), respectively. These distributions differ slightly in passing from subunit $\boldsymbol{A}$ to subunit $\boldsymbol{I}$, confirming the asymmetry in the two subunits.

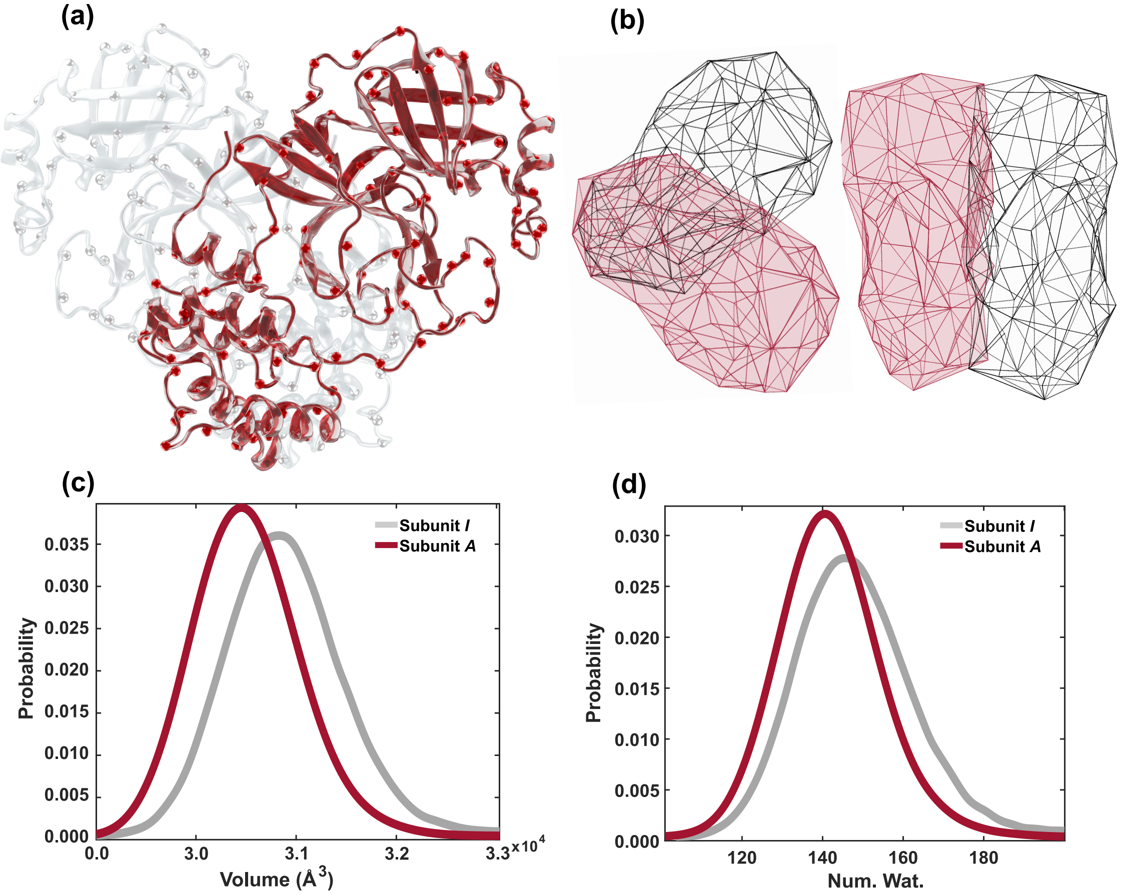


Figure S-2: (a) SC-2 M$^{pro}$ subunit $\boldsymbol{A}$ (red) and $\boldsymbol{I}$ (grey). The $\alpha$-Carbon atoms of the residues located on the protein outer-surface are represented by small spheres. The same conformation as that of Fig. S-1 is shown. (b) Convex hull surfaces built from the $\alpha$-Carbon atoms shown in (a) for each subunit. Distribution of (c) each subunit's volume inside the convex hull and of (d) the number of water molecules that they contain, estimated over all of the DESRES trajectory.

Table S-1: The six water molecules with the lowest Debye-Waller factor of the SC-2 M$^{pro}$ X-ray structure by Owen et al. [2] (PDB entry 6Y84). All other water molecules have a Debye-Waller factor > 15. The data are identical for both subunits.

| **Index of water molecules from PDB entry 6Y84** | **Debye Waller factor** |
|---|---|
| **2639** | 14.76 |
| **2664** | 12.60 |
| **2583** | 14.96 |
| **2721** | 14.44 |
| **2679** | 12.16 |
| **2778** | 11.47 |

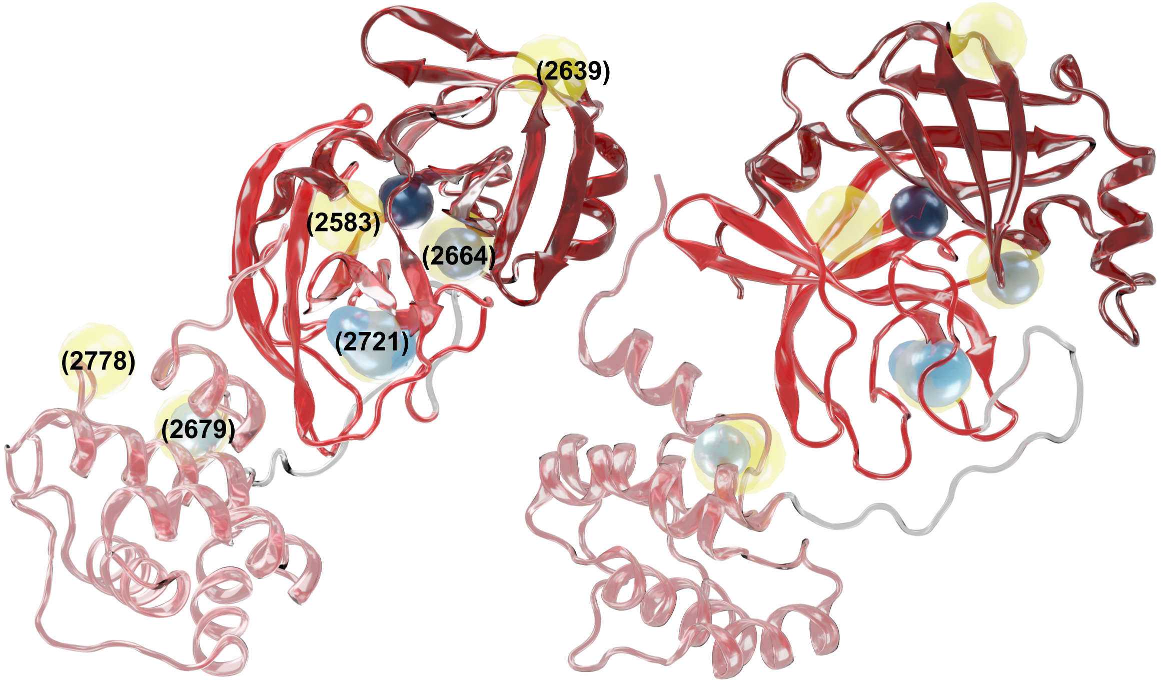


Figure S-3: Water molecules with a low Debye-Waller factors reported in Tab. S-1 (yellow spheres), along with *long lived water molecules* (blue spheres) inside one subunit of the SC-2 M$^{pro}$ from two different orientations. Darker blue spheres indicate longer lifetimes. Only one subunit is shown because of the symmetry in the long lived water molecules position (see the main text for more details).

### S2 Mobility of the water molecules inside the protein

#### S2.1 Low mobility

Fig. S-3 shows the *long-lived water molecules* (see also Fig. 2 (b) in the main text) inside the convex hull surface of subunit *A*, along with 6 water molecules with the lowest Debye-Waller factors in the X-ray structure (Tab. S-1) (yellow and blue spheres, respectively). Three of the *long-lived water molecules* overlap with water molecules 2664, 2721, 2679 in the PDB file. One is in proximity of water 2583. Water molecules 2639 and 2778 are located on the protein's surface in the solid state structure and, not unexpectedly, are mobile in aqueous solution.

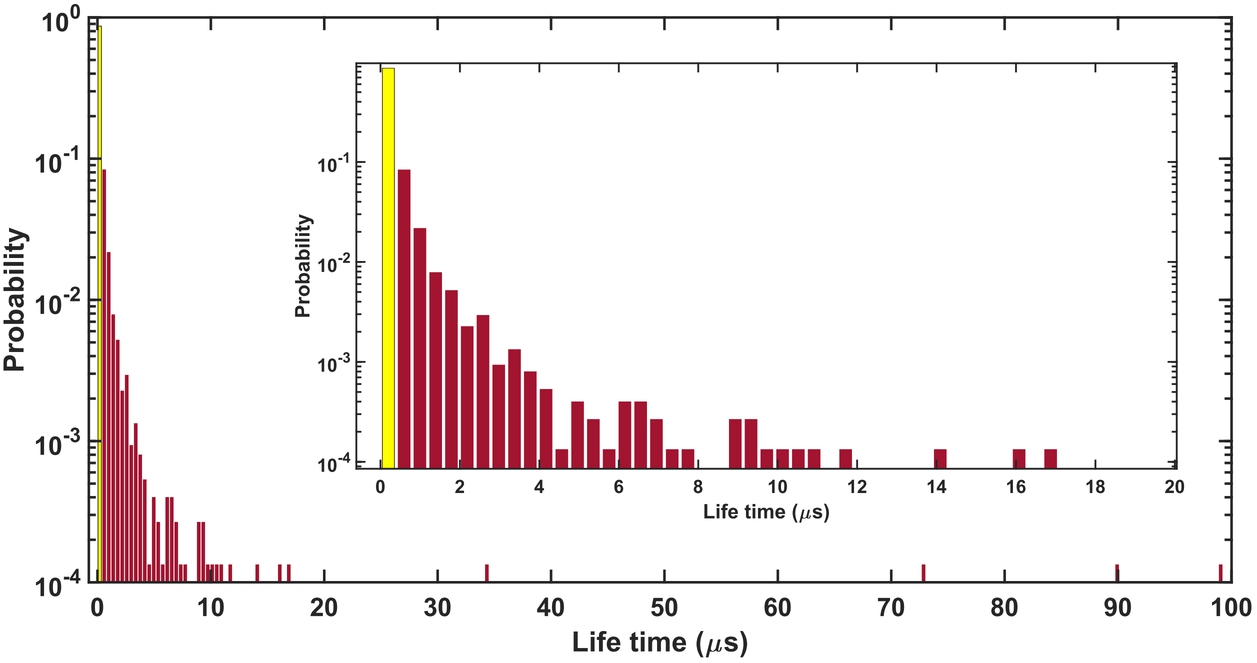


Figure S-4: Distribution (on a logarithmic scale) of water molecules with a residence time inside the convex hulls greater than 100 ns and 0.5 $\mu$s (red and yellow bars, respectively). The inset shows the region of residence time less than 20 $\mu$s.

#### S2.2 All ranges of mobility

Fig. S-4 shows the lifetime distribution of the water molecules whose residence times inside the convex hull surfaces are at least 100 ns in the DESRES simulation. The inset shows the distribution of water molecules within the 100 ns - 20 $\mu$s window. About 87 % of the considered water molecules have a lifetime in the 100-500 ns range. Conversely, around 12.5 % of them feature a lifetime between between 0.5 $\mu$s and 10 $\mu$s (*mid-lived water molecule*, while only ~0.5 % of water molecules have a lifetime between 10 $\mu$s and 99 $\mu$s (*long-lived water molecules*).

### S3 Hydration sites

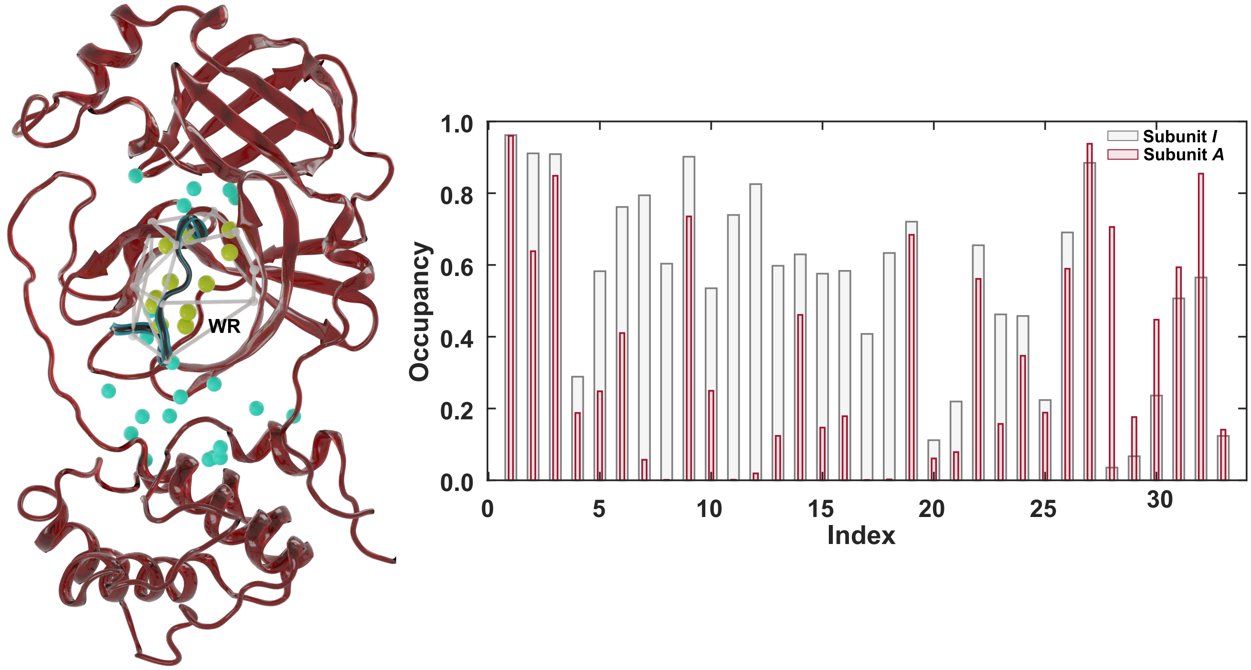


Figure S-5: Hydration sites (HSs) in the SC-2 M$^{pro}$ protein, as emerging from an analysis of the DESRES trajectory. Left hand side: Subunit ***A*** (red) with its 33 (HSs), represented as spheres. The water reservoir region (WR), defined in section S4, is shown in grey lines and its HSs inside in yellow. The m-shaped loop is coloured in blue. Right hand side: The occupancy of each HS in the active ***A*** and the inactive ***I*** subunits in the DESRES trajectory. A value of 1 correspond to an HS that is always occupied by a water molecule throughout the trajectory.

To identify the hydration sites (HSs), namely the spots where water molecules are mostly located, we select the water molecules with a residency time inside the convex hulls of at least 500 ns (see Sec. S2.2). The HSs are then identified by dividing the space inside the convex hulls in a grid and use a k-means cluster analysis [3]. We identify 33 HSs for each subunit (see Fig. S-5). Next, we assign the same label to an HS if it appears in a symmetry-equivalent positions in both subunits. Finally, we calculate the water occupancy of the HSs on a sphere of radius 1.5 Å centred on them, along the entire DESRES trajectory. If a water molecule appears inside such a sphere at all times, the occupancy is 1. The occupancy of most HSs in subunit ***I*** turns out to be significantly higher than that in subunit ***A*** (Fig. S-5).

### S4 Volume and number of water molecules contained in S1 binding pocket.

In this section, we investigate the change in water content associated with the symmetry breaking transition observed in the DESRES trajectory. Rather than considering the entire protein, we focus therefore on the region around the S1 binding pocket, that we call *water reservoir* (WR) hereafter. In fact, the WR (i) contains a large number of HSs (see Fig. S-5) and (ii) it is the region where the transition begins. Let us define a convex hull enclosing the WR and evaluate its volume and water molecules content along the

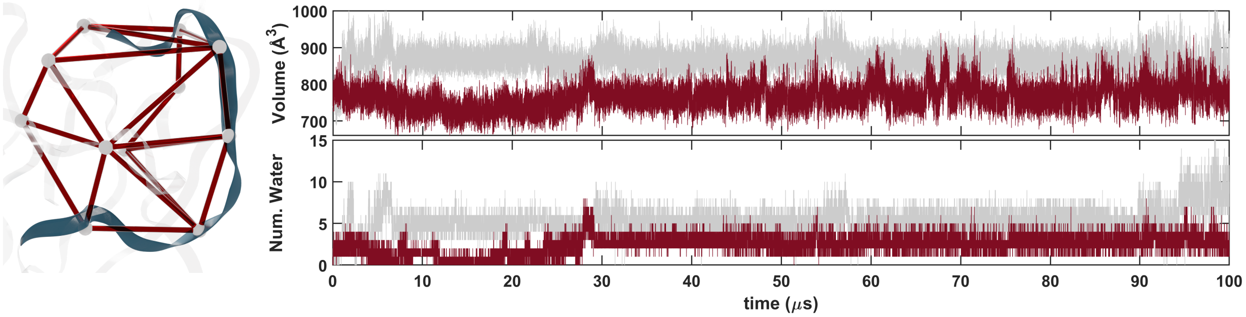


Figure S-6: Left hand side: Cartoon representation of the water reservoir (WR, see text) around the S1 binding pocket shown in Fig. S-5, along with the convex hull enclosing it. As in Fig. S-5, the m-shaped loop is coloured in blue. Right hand side: WR volume and number of water molecules for subunit ***A*** (red) and subunit ***I*** (grey) plotted as a function of simulated time in the DESRES trajectory.

DESRES trajectory[1]. Fig. S-6 shows that, in subunit ***I***, these quantities start from a similar value than those of subunit ***A***. Then, after about 1 $\mu$s, they become larger than those of subunit ***A***. This abrupt change is associated with the break of symmetry in the homodimer.

[1] The same calculation for the entire subunit (Sec. S1) does not show a significant change of these quantities as the signal to noise ratio associated with a convex hull covering all the subunit is much lower.

#### S5 Symmetry of the two subunits

The symmetry between subunits across a variety of PDB structures is estimated here in terms of the root-mean-square deviation (RMSD) values between the backbone atoms. We use the VMD program [4] (see Tab. S-2). The RMSD values for the conformations of the DESRES trajectory fall in a range between ∼0.5 and ∼2.7 Å.

Table S-2: The root mean square deviation (RMSD) (Å) between the backbone atoms of two subunits in X-ray structures of SC- and SC-2 M$^{pro}$ proteins. Structures with a ligand bound to the S1 binding pocket of both subunits are indicated with a *.

| | PDB ID | residues | RMSD |
|---|---|---|---|
| **SARS-CoV** | 2bx3 | 3 - 300 | 0.00 |
| | 1p9s* | 5 - 290 | 0.35 |
| | 1uk2 | 5 - 290 | 0.74 |
| | 1uk3 | 5 - 290 | 0.79 |
| | 1uj1 | 5 - 290 | 2.65 |
| **SARS-CoV-2** | 6y84 | 5 - 300 | 0.00 |
| | 6y2g* | 5 - 300 | 0.21 |
| | 7lkd | 5 - 300 | 0.25 |
| | 6xqt* | 5 - 300 | 0.31 |
| | 7joy | 5 - 300 | 0.91 |
| | 7lmc | 5 - 300 | 1.03 |

#### S6 Details of the simulations carried out in this work

We take as initial configuration of SC-2 M$^{pro}$ a snapshot of the DESRES simulation after 60.2 microseconds [1]. The system is enclosed in a cubic box of 115.6 Å, 115.6 Å, 115.6 Å sides with a total number of 49,716 water molecules. Periodic boundary conditions are applied. The protein, the ions and water are described by the Amber99sb-ildn [5] and TIP4P-D [6] force fields, respectively. A Ewald [7] summation is applied for the long-range electrostatic interactions with a cutoff of 12 Å for both the real part of the Coulomb interactions and the van der Waals interactions.

Constant pressure and temperature simulations are achieved by using the Parrinello–Rahman barostat [8] and the stochastic velocity rescaling thermostat [9]. For both thermostat and barostat, the coupling constant is 1 ps. All bonds involving hydrogen atoms are constrained using the LINCS algorithm [9]. The SETTLE algorithm is used for water [10].

The calculations are performed with the MD software GROMACS-2020.4 [11]. For enhanced sampling, we use a custom version of the PLUMED 2.7 plugin [12] where we include the Pytorch library 1.4 [13].

#### S7 Plain molecular dynamics

The system is equilibrated for 10 ns, first in the NVT ensemble and then in the NPT ensemble. The production run is in the NPT ensemble at 300 K and 1 atm. We integrate Newton's equations of motions using the leapfrog algorithm with a time step of 2 fs.

#### S8 Enhanced sampling simulations

We perform biased simulations using the on-the-fly probability enhanced sampling (OPES) [14] technique, a recent evolution of Well-Tempered Metadynamics [15, 16], that is able to calculate the exact free energy of a process as a function of suitable collective variables [17]. The method builds a time-dependent bias from an on-the-fly estimate of the probability distribution. This estimate is built by depositing Gaussian kernels at fixed time intervals. OPES determines an intense initial exploration of phase space that is followed by a quasi-static regime where the bias changes very little and a robust convergence is reached. Phase space exploration is limited by the activation barrier parameter that effectively sets a hard limit on how much energy above the free energy minimum the method can explore. For this work, it is crucial to identify CVs that are able to distinguish active and inactive subunits in SC-2 $M^{pro}$ and drive transitions between them.

Here, inspired by the work of some of us [18], we use the recently developed machine learning Deep-LDA method [19]. Deep-LDA is a dimensionality reduction technique which takes the thermal fluctuations of a set of descriptors in two states and uses them to train a neural network (NN). The latter produces a CV that is meant to distinguish the two states. Our NN architecture consists of a sequence of layers with $18, 14, 12, 6, 4$ nodes both CVs, with the rectified linear unit as activation function. The two states that we use for training are the active and inactive subunits of the homodimer in the DESRES trajectory between 30 $\mu$s to 90 $\mu$s.

We apply Deep-LDA to generate two different CVs. One CV, $s_c$, focuses on the protein structural changes between the active and the inactive subunit, the other one, $s_w$, is based on the water distribution.

To identify $s_c$, we combine in total 26 descriptors. These descriptors are of three types:

1) The coordination between four HSs and $\alpha$-Carbon atom of residues 135-146 (see panel Fig. S-7 (a)).

2) The coordination number between inter-subunit residues (see panel Fig. S-7 (b)):
[$O_{ASN142}$, $NH_{GLY2}$], [$O_{PHE140}$, $NH_{PHE3}$], [$NH_{PHE140}$, $O_{PHE3}$], [$O_{SER139}$, $NH_{ARG4}$], [$O_{LYS137}$, $NH_{ARG4}$], [$N_{SER1}$,$O_{GLU166}$], [$S_{MET6}$,$C_{TYR126}$] and [$N_{ARG298}$,$O_{MET6}$].

3) The coordination number between intra-subunit residues (see panel Fig. S-7 (c)):
[$C_{HIS141}$, $S_{CYS145}$], [$O_{GLU166}$, $C_{HIS172}$], [$O_{GLU166}$, $C_{PHE140}$], [$O_{GLU166}$, $C_{HIS172}$], [$O_{HIS164}$, $N_{HIS163}$], [$N_{HIS163}$, $N_{HIS172}$], [$N_{HIS163}$, $N_{HIS172}$], [$N_{HIS163}$,$O_{GLU166}$], [$N_{HIS41}$,$O_{HIS164}$], [$O_{GLU166}$,$N_{SER144}$], [$O_{GLU166}$,$N_{GLY143}$], [$C_{ALA116}$,$C_{PHE140}$], [$S_{CYS145}$,$N_{HIS172}$], and [$S_{PHE140}$,$_{TRY126}$].

Fig. S-7 shows the trajectories of a number of descriptors in ***A*** and ***I*** for each of the above mentioned steps.

For the CV $s_w$, we use the coordination number of water around 33 HSs with the functional form :

$$\mathrm{d}_i^0 = \sum_j^{r_{ij}<r_{\mathrm{NL}}} \frac{1-\left(\frac{r_{ij}}{r_0}\right)^n}{1-\left(\frac{r_{ij}}{r_0}\right)^m} \tag{S-1}$$

where $r_{ij}$ measures the distance between the centre of HS $i$ and the water oxygen atom $j$ within a sphere of radius $r_{\mathrm{NL}} = 10$ Å and $r_0 = 2.5$ Å. We set $n = 2$ and $m = 6$.

We use OPES to explore the possibility of a flip between subunits in the asymmetric homodimer. We bias the $s_c$ CVs of both subunits starting from an equilibrated asymmetric homodimer configuration from the DESRES trajectory. We use a deposition pace of 2 ps and employ different activation barriers. If the barrier parameter is too low, no transition is observed as the system cannot surmount the barrier pertaining to the flip event. By observing the CV trajectories for different barriers, we can get a rough estimate of the real activation barrier for the subunit flip process. The lowest barrier that allows a flipping event can be considered our barrier estimate.

Fig. S-8 shows the value of $s_c$ as a function of time in simulations with different OPES activation barriers, with the initially active subunit in red and the initially inactive one in grey. All applied activation barriers could trigger a flip event on the initially active subunit, changing it to inactive. On the other hand, the initially inactive subunit remains largely unchanged for barriers lower than 125 kJ/mol. Only for an activation barrier between 125 kJ/mol to 150 kJ/mol (see panel (c,d) of Fig. S-8) we observe a flip of the initially inactive subunit into active, completing the subunit exchange. Such a high barrier confirms the high stability of the asymmetric homodimer state, making the subunit flip an extremely rare event at room temperature.

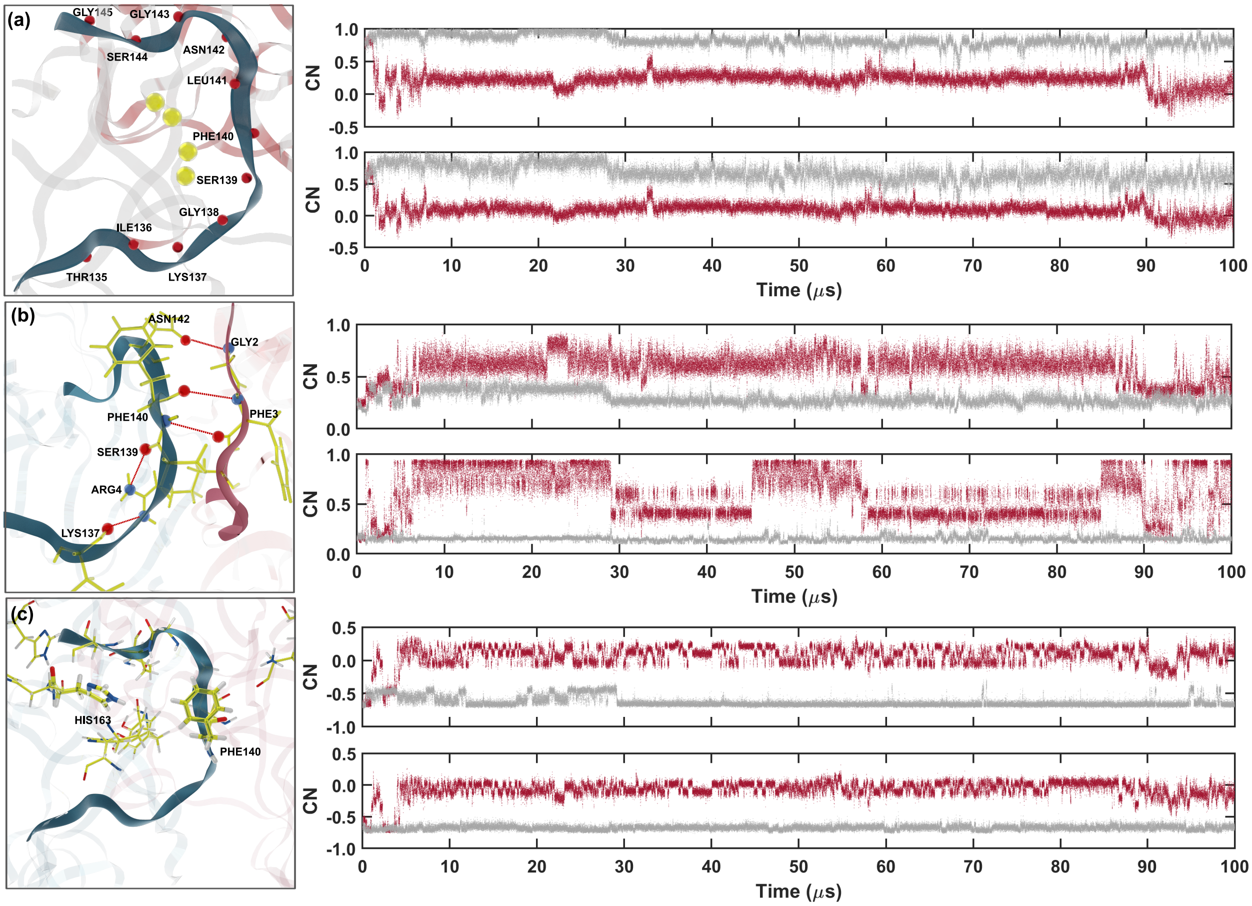


Figure S-7: Left: the WR region (see text) with a focus on different residues and their interactions according to type (1) (panel a), (2) (panel b) and (3) (panel c), described in section S8. Right: representative descriptors trajectories corresponding to type (1) (panel a), (2) (panel b) and (3) (panel c). The quantities are plotted as a function of simulated time in subunit $\boldsymbol{I}$ (gray) and $\boldsymbol{A}$ (red).

During the simulations, we monitor the presence of water inside the subunits by checking the trajectory of $s_{\mathrm{w}}$. Fig. S-9 shows the trajectory of $s_{\mathrm{c}}$ and $s_{\mathrm{w}}$ CVs in both subunits in a simulation with an initial barrier of 400 kJ/mol. The transition events in $s_{\mathrm{c}}$ are accompanied by large fluctuations and transitions of $s_{\mathrm{w}}$, indicating that water is not the slowest degree of freedom of the event we are studying.

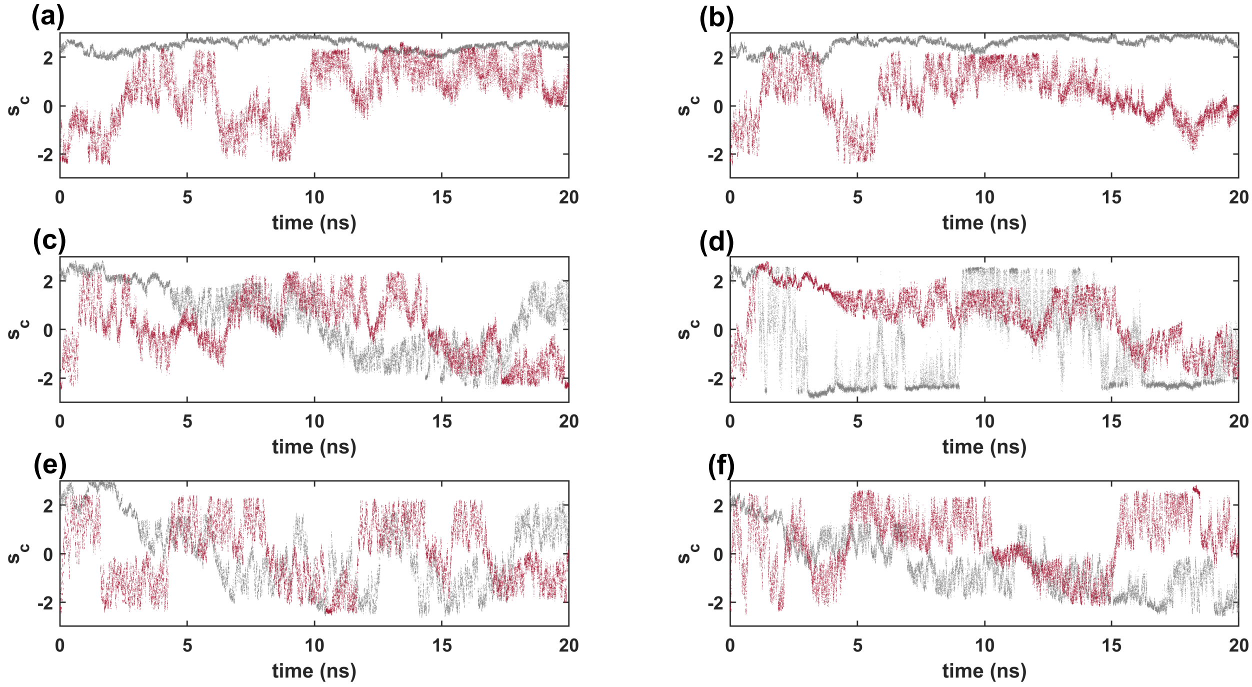


Figure S-8: The value of $s_c$ in simulations with an OPES barrier guess equal to (a) 100 kJ/mol, (b) 125 kJ/mol, (c) 150 kJ/mol, (d) 200 kJ/mol, (e) 300 kJ/mol, and (f) 400 kJ/mol. The initially active subunit is shown in red and the initially inactive one in grey.

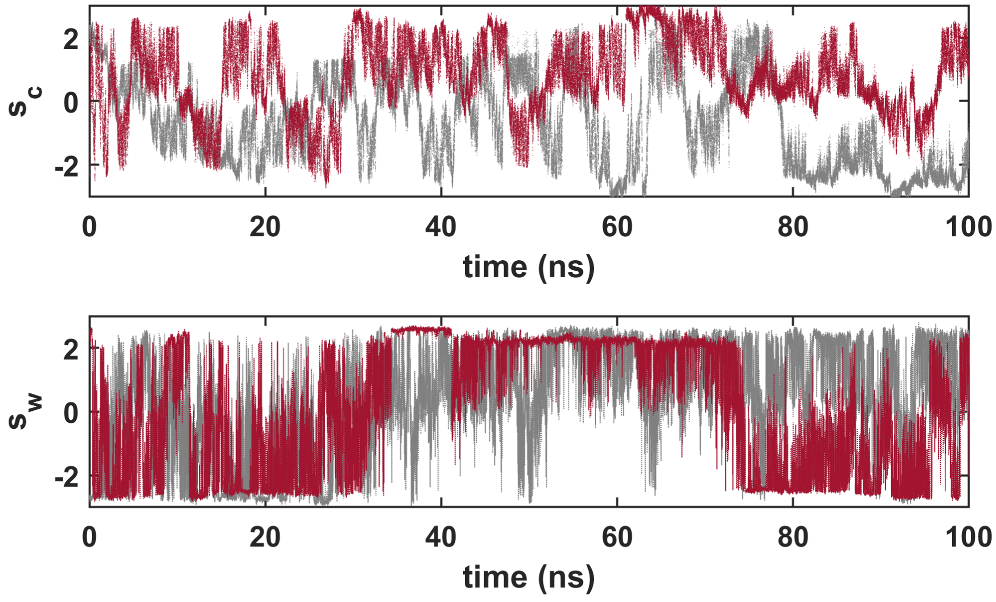


Figure S-9: The values of $s_c$ and $s_w$ in a 100 ns simulation with an activation barrier equal to 400 kJ/mol. The initially active subunit is shown in red and the initially inactive one in grey.
